# Concurrent neural representations of the current and forthcoming movement plans in the frontal eye field during a saccade sequence

**DOI:** 10.1101/390658

**Authors:** Debaleena Basu, Aditya Murthy

## Abstract

We use sequences of saccadic eye movements to continually explore our visual environments. Previous studies have established that saccades in a sequence may be programmed in parallel by the oculomotor system. In this study, we tested the neural correlates of parallel programming of saccade sequences in the frontal eye field (FEF), using single-unit electrophysiological recordings from macaques performing a double-step saccade task. Neurons in the FEF range from visual neurons instantiating target selection, to movement neurons which prepare a saccadic response towards the selected target. The question of whether the FEF movement neurons undertake concurrent processing of multiple goals or saccade plans is yet unresolved. We show that when a peripheral target is foveated by a sequence of two saccades, FEF movement activity for the second saccade can be initiated whilst the first is still underway. Moreover, the onset of the movement activity varied parametrically with the behaviorally measured time available for parallel programming. Finally, the concurrent activity was specific for the final remapped motor vector connecting the first and the second targets and not the goal of the second saccade. In contrast, the upstream FEF visual-related neurons showed concurrent activity related to the goal of the second saccade, but not the remapped vector connecting the first and the second targets. Taken together, the results indicate that movement neurons, although located terminally in the FEF visual-motor spectrum, can accomplish concurrent processing of multiple saccade plans, leading to rapid execution of saccade sequences.

## Introduction

Saccadic eye movements serve to rapidly shift the fovea to various parts of a surrounding scene for visual inspection. Many day-to-day tasks necessitate the execution of a meaningful sequence of multiple saccades. The simple act of reading a letter, for example, requires planning and execution of saccade sequences aimed at scanning the page from top to bottom. The question thus arises: how are multiple saccade plans linked into a coherent sequence?

At the level of execution, saccadic behavior is inherently serial: a forthcoming saccadic eye movement can only happen after the current one is completed. At the level of neural programming as well, a sequence of saccades may be planned in a purely serial fashion, with the sequence emerging as a simple concatenation of multiple single saccade plans. Alternatively, the oculomotor system may be able to operate in parallel, wherein programming of an upcoming saccade may begin before the conclusion of the ongoing saccade. Various lines of behavioral evidence have postulated the possibility of parallel programming of saccade sequences by the oculomotor system (Becker and Jürgens 1979; Minken et al. 1993; McPeek et al. 2000; McPeek 2003; Ray et al. 2004; Sharika et al. 2008; Bhutani et al. 2012, 2013; Wu et al. 2013).

Motivated by established behavioral data, we explored the underlying neural correlates of parallel programming of saccade plans, in the frontal eye field (FEF), an oculomotor area critical for saccade programming (Schall et al. 2002a, 2013; Sendhilnathan et al. 2017). FEF neurons show functional differences based on whether the neuronal activity is evoked in response to stimulus appearance (visual neurons) and/or whether the activity leads to saccade initiation (visuo-movement neurons/movement neurons; Bruce and Goldberg 1985; Hanes and Schall 1996). FEF visual neurons have been shown to instantiate the stage of target selection, wherein the target for the next saccade is located from the current visual scene (Thompson et al. 1996). Target selection is delinked from the saccadic response as it can occur even when saccades are stopped (Juan et al. 2004). Studies in which the saccade goal was incongruent with the visual stimulus, or where the visual and motor phases are separated by a delay, have shown that FEF visual responses generally encode the location of the visual stimulus, whereas FEF movement responses are tuned to the actual saccade goal to which the saccade has to be executed (Everling and Munoz 2000; Sato and Schall 2003; Takeda and Funahashi 2004; Sajad et al. 2016). Additionally, visual cell activity does not predict reaction times in simple oculomotor tasks (Hanes et al. 1998).

In view of the existing results indicating a separation between the perceptual and motor roles of the FEF (Juan et al. 2004; Costello et al. 2013), it is essential that direct evidence for concurrent saccade programming relies on analyzing the activity of movement-related cells of the FEF, which has been shown to predict behavioral mesures such as reaction times. Our motivation to focus on the FEF movement cells also follows from the fact that the bulk of the evidence for parallel processing comes from temporal indicators of behavior such as intersaccadic intervals (ISI).

Parallel programming of a two-saccade sequence is made complicated by the fact that it involves a visuo-motor transformation of the retinal vector encoding the second target location (from *retinal vector 2* to *saccade vector 2*\* in **Fig. 6A**), in order to predictively compensate for the retinal displacement produced by the first saccade, and accurately generate the second saccade rapidly from the first target. Any concurrent activity in the FEF movement neurons, can, thus either encode the retinal vector of the second saccade goal, or the remapped second saccade vector. Using midway saccades in an adaptation task, Bhutani et al. (2017) recently showed that the remapped second saccade vector can be planned before the first saccade, indicating a similar representation in the brain.

Till date, while FEF visual responses have been looked in the context of saccade sequencing (Tian et al. 2000; Phillips and Segraves 2010), there has not been a comprehensive effort to understand the contribution of the execution-linked, downstream FEF movement neurons in sequential saccade processing. We aimed to determine if, at the level of FEF movement neurons, there was a neural correlate for parallel programming of a second saccade whilst the first was still underway. This notion was tested using a modified version of the classical double-step task (Westheimer 1954), the FOLLOW task. Based on previous findings of concurrent activity in the superior colliculus (McPeek and Keller 2002; Port and Wurtz 2003; Shen and Pare 2014), an important oculomotor area downstream of FEF, our speculation was that FEF movement neurons will show activity encoding the second saccade goal, even as the first saccade is being executed. Based on behavioral evidence from midway saccades (Bhutani et al. 2017), we also hypothesized that the concurrent representation would be of the remapped motor vector of the second saccade, allowing rapid initiation of the consequent movement and leading to shorter ISIs, corroborating with previously observed behavior.

## Materials and Methods

### Animals

The objectives of this study were investigated by analyzing behavioral and neural data collected from two adult monkeys (**J**, male *Macaca radiata*; age 9 years, weighing 5.5 kg, bred in-house, and **G**, female *Macaca mulatta*; age 11 years, weighing 3.8 kg, acquired from the National Brain Research Centre, India). The animals were cared for in accordance with the animal ethics guidelines of the Committee for the Purpose of Control and Supervision of Experiments on Animals (CPCSEA), Government of India, and the Institutional Animal Ethics Committee (IAEC) of the Indian Institute of Science. All protocols followed were in accordance with the guidelines set by the National Institutes of Health, USA.

### Surgical Procedures

Two sterile surgeries were performed on each monkey to for fixing of a titanium head post and making a craniotomy over the FEF, respectively. The positioning of the Cilux plastic recording chamber (Crist instruments, USA) was guided by MR-images (Philips Achieva, 3T for monkey ’G’ and Philips Achieva, 1.5 T for monkey ’J’), and stereotaxic coordinates. During the surgery, the chamber placement was further confirmed by identifying the arcuate sulcus through the exposed dura mater. Both the monkeys had chambers placed over the right FEF. Behavioral training was resumed only after complete recovery from the surgeries.

### Experimental tasks

The monkeys were trained on two oculomotor tasks for the purpose of the experimental studies. In the memory-guided (MG) saccade task, each trial started with a red fixation point (0.6° × 0.6°) appearing at the center of a screen. The monkeys were required to fixate on the central spot for a variable amount of time (~300 ms), following which a gray target stimulus (1° × 1°) appeared at a peripheral location. The target could appear at any of the eight possible locations on an imaginary circle of radius 12°, centered on the fixation point. Post-appearance, the target disappeared after 100 ms. The monkeys were required to continue fixation after the target flash for an additional delay period (~1000 ms). After the delay period, the fixation spot disappeared, cueing the monkeys to make a single saccade to the memorized location of the target. Electronic error windows of ± 4°-6° drawn around the fixation point and target, were used to assess the online performance. A liquid reward was delivered after every correct response. The delay period separated the stimulus-related (visual) and saccade-related (motor) epochs, allowing classification of FEF neurons. The MG task was also used to determine the response field of the neurons.

The FOLLOW task (**Fig. 1**) is a modified version of the classical double-step task (Westheimer 1954; Wheeless et al. 1966; Becker and Jürgens 1979), which was designed to generate a sequence of two purposeful, visually-guided saccades. The task consisted of two types of trials that were randomly interleaved: *no-step* trials (30%), and *step* trials (70%). Each trial started with the appearance of a white central fixation point (0.6° × 0.6°). In *no-step* trials, following fixation for a variable duration (~300 ms), a peripheral green target (1° × 1°) appeared in one of six possible locations (see next paragraph) at an eccentricity of 12° centered at the fixation point. The appearance of the target was accompanied by extinguishing of the fixation point, thus signaling the monkeys to make an immediate saccade to the target. Two targets appeared in *step* trials. The initial target was identical to the single no-step target. A red final target (1° × 1°) appeared after the first target following a variable time delay, referred to as the target step delay (TSD). Three TSDs were used: 16 ms, 83 ms, 150 ms. The correct response in the step trials was to make a sequence of two saccades: from fixation point to the first target, and from the first target to the second target. Electronic error windows of ± 4°-6° drawn around the fixation point and targets were used to evaluate online performance. The windows were kept large to accommodate variability in the larger second saccades, and encourage rapid execution as opposed to absolute end-point accuracy. Correct responses were rewarded with liquid reward.

**Figure 1.**
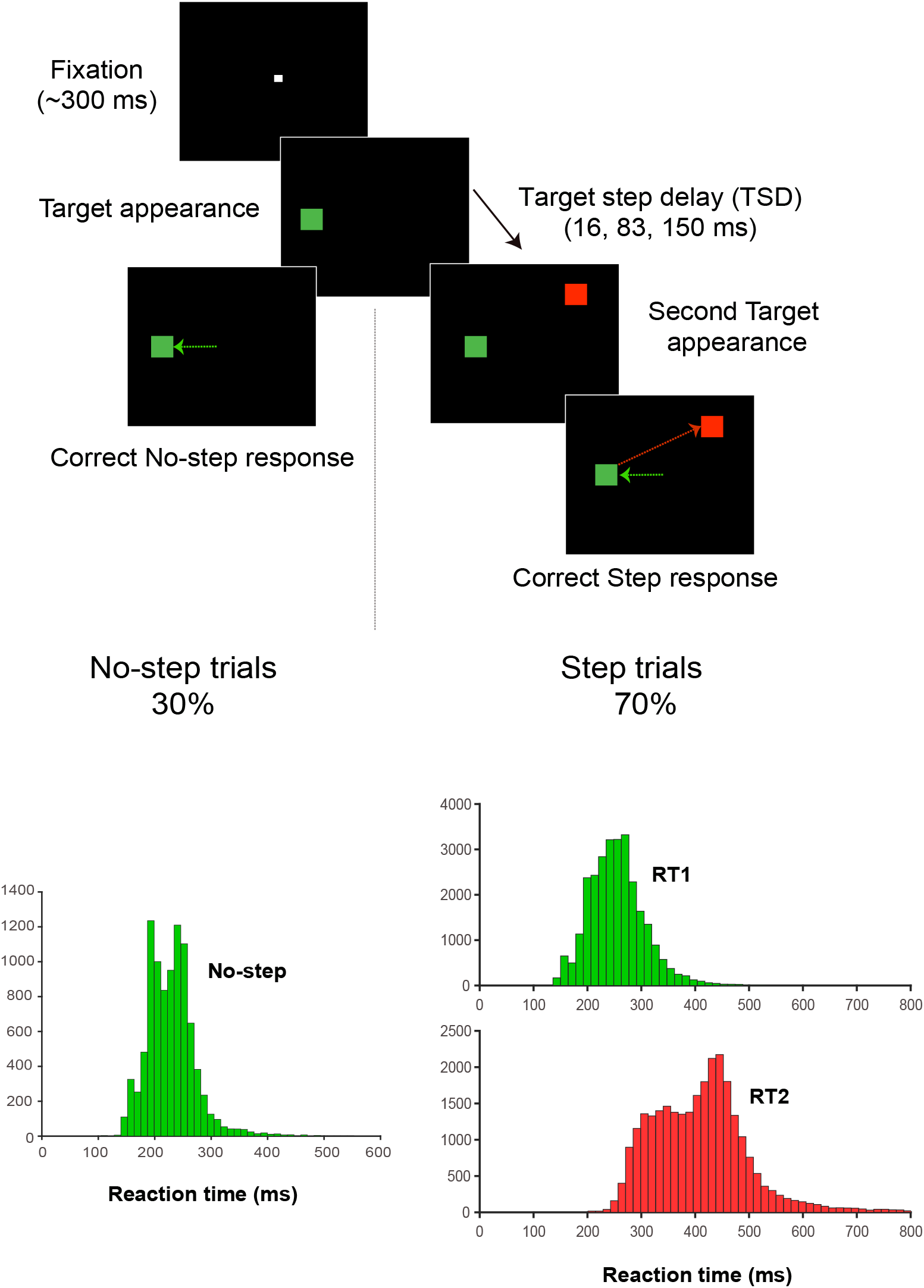
FOLLOW task paradigm. Trials in the sequential saccade FOLLOW task started with central fixation, following which a green target was presented at any 1 of 6 possible locations (see Materials and Methods). In no-step trials (30%), the monkeys were required to make a single saccade to the target for juice reward. In randomly interleaved step trials (70%), a second red target appeared after a variable target step delay (TSD) and the monkeys had to execute a sequence of two saccades to the targets. TSD varied randomly across step trials. The bottom panel shows the reaction time distributions for no-step trials (*left*), and the first (RT1) and second (RT2) saccades of the step trials (*right*).

While the MG task had 8 possible target locations, a restricted set of target positions and steps were used during neurophysiological recording sessions to maximize collection of relevant FOLLOW task data. After response field (RF) identification in the MG task, typically three target locations centered on the RF were considered to be ‘inside-RF’ positions and the three positions diametrically opposite to them were considered to be out of the response field or ‘outside-RF’ positions. The targets in no-step trials and the first targets in step trials could appear in any one of the 6 inside-RF *and* outside-RF positions. However, the second target in step trials could only appear in one of the three positions diametrically opposite to the position of the first target. Thus, the second target could either step into or out of the response field but never within or adjacent to it. ***RFin trials*** refer to trials in which the second target stepped or appeared inside the RF while the first target appeared outside of RF. Therefore, the neural response would mainly encode the second target or second saccade. ***RFout trials*** were those in which the second target appeared outside of RF after a first target appeared within it. Thus, the neural responses would mainly be for the first target or first saccade. *RFin* trials constitute the main trial set of interest for this study.

The step trials had 18 possible target-step arrangements: 6 possible target 1 locations × 3 possible target 2 locations for each first target. The targets could appear in any of the possible locations with equal probability. The hemifield separation maintained between the first and second targets also ensured that the angular separation between both the targets was least 90°, reducing the chances of averaged or curved saccades.

### Recording setup and procedures

The task paradigms were designed and controlled using TEMPO software (Reflective computing, St. Louis, MO, USA). Task stimuli were displayed using VIDEOSYNC software (Reflective computing) working in tandem with TEMPO. All stimuli were presented on a Sony Bravia LCD monitor (42 inches, 60 Hz refresh rate; 640 × 480 resolution) placed 57 cm from the monkeys so that 1° of visual angle would correspond to 1 cm on the monitor. All the experiments were conducted in a dark room with head-restrained monkeys, sitting in their respective primate chairs. A spout, put in the monkey’s mouth, delivered the liquid reward. Eye position was sampled at 240 Hz with a monocular infrared pupil tracker (ISCAN, Woburn, MA USA) that interfaced with the TEMPO software in real-time. All task parameters were saved in the TEMPO system with a sampling rate of ~1000 Hz.

Single-unit recordings were done using epoxy-insulated tungsten electrodes (FHC, Bowdoin, ME, USA, impedance 2 - 4 MΩ) having a diameter of 0.2 mm. The electrodes were advanced into the brain using a hydraulic microdrive setup (FHC) and unit activity was characterized using the MG task at intervals of 50–100 μm in depth. Neuronal data was sampled and stored at 30,000 Hz by the Cerebus data acquisition system (Blackrock Microsystems, Salt Lake City, UT, USA). A unit was confirmed to be an FEF neuron if the activity matched that of established FEF cell activity types (Bruce and Goldberg 1985). The position evoking the highest activity in the MG task was chosen as the RF of the unit. Once the neuron type and RF was well characterized in the MG task (~ 100-200 trials), the FOLLOW task was started and continued till sufficient data (~700 trials) was collected.

### Data analysis

Neural data was sorted into individual units offline using time-amplitude algorithms provided with the Cerebus Central Suite (Blackrock Microsystems). Saccade onset and offset times were detected from the eye position data using a 30°/s velocity criterion. All analyses, after spike sorting, were done using custom-made scripts written in MATLAB (MathWorks, Natick, MA, USA).

Only correct trials, wherein saccade parameters like direction, and order, were accurate, and saccadic end-points were well within the electronic windows bounding the target stimuli, were analyzed. For the FOLLOW task, trials with saccadic latency shorter than 80 ms were considered anticipatory saccades and rejected. Delayed saccades crossing the maximum permissible saccade latency limit (600 ms for *no-step* trials and first saccade of *step* trials; 900 ms for second saccade of *step* trials) were also excluded from analysis. A minimum of 8 trials per condition was set as the criteria for performing any analysis. Units which could not be isolated clearly or which did not show spatially selective, task-related responses were discarded. After applying the exclusion criteria, a total of 87 neurons were selected for further analysis and contributed to the results of this study.

Spike density functions (SDFs) for neural data were calculated by convolving the averaged spike train with a filter that resembled an excitatory post-synaptic potential (EPSP), having a combination of growth and decay exponential functions (Thompson and Schall, 1997). The time constants of the rapid growth phase (τ_g_ = 1 ms) and the slower decay phase (τ_d_ = 20 ms) were matched to values obtained from excitatory synapses and thus, were physiologically relevant.

FEF neurons were classified as visual, visuomovement, or movement by evaluating the activity during the memory-guided saccade task. Visual activity (VA) was defined as the mean firing rate in the visual epoch (100-180 ms after target onset) above the mean spontaneous firing rate measured from the baseline epoch (300-100 ms preceding target onset). Movement activity (MA) was the mean firing rate in the movement epoch (80 ms preceding saccade onset) above the baseline firing rate. *Visual neurons* significantly increased their firing rate compared to baseline in the visual epoch, post-stimulus presentation, but showed no significant activity build-up in the movement epoch. *Movement neurons* showed ramping up of activity prior to saccades in the movement epoch, but no stimulus-related activity. *Visuomovement neurons* showed heightened activity in both the visual and movement epochs.

To quantify the proportions of visual- and movement-related activity, a *visuo-motor index* (VMI) was calculated for each neuron (Murthy et al. 2007).

VMI was calculated as: VMI = (VA-MA) / (VA+MA)

The VMI can range from +1 to −1. Visual neurons with comparatively higher VA yielded positive VMIs, whereas movement neurons with higher MA had negative VMIs. Visuomovement neurons had intermediate VMIs.

The onset of second saccade related activity or the **Neural selection time (NST)** for each neuron was defined as the time point when differential activity between *RFin* and no-step-out trials of the FOLLOW task, first exceeded 2 standard deviations (SDs) above baseline activity (100 ms period prior to onset of first target), provided the differential activity ultimately reached 4 SDs and was maintained above 2 SDs for at least 20 ms.

The divergence point between the *RFin* and no-step-in trials (**Fig. 6**) for each movement neuron was calculated similarly for latency-matched trials (PPT in RFin trials was matched to RT of no-step trials), except that the difference histogram (RFin activity – no-step activity) could either exceed or fall below the aforementioned limits.

For calculating the population divergence point averaging across the visual, visuomovement, and movement neurons, (**Fig. 7**), a similar process was used, but with a more stringent criterion: after the putative divergence point, the differential activity had to be maintained above or below 2 SDs of the baseline activity for at least 50 ms, within which it has to reach ±6 SDs of the baseline activity. The stringency was increased as this was a population–based measure, and thus expected to have lesser signal variation as opposed to activity of single neurons.

### Statistical testing

Single sample analysis was done using a two-sided Wilcoxon signed-rank test. For group comparisons with a single factor, one-way ANOVA (analysis of variance) was used. The trials of various TSD groups were randomly interleaved in the experimental design and thus were considered to be independent observations. Normality in each dataset was tested using the Lilliefors test. For non-normal data sets, the non-parametric version of ANOVA, the Kruskal-Wallis test was used. All the results are presented as mean (± standard error of mean, SEM) and all tests are performed at a significance level of α = 0.05 unless otherwise mentioned.

## Results

One *Macaca radiata* (J) and one *Macaca mulatta* (G) performed a sequential saccade ‘FOLLOW’ task (**Fig. 1**; see Materials and Methods for details), where, in about 70% of the trials (*step trials*) they had to execute a sequence of visually-guided saccades to two target stimuli in their order of appearance. In about 30% of the trials (*no-step trials*), a single target was presented to which a single saccade had to be made. Step and no-step trials were randomly interleaved. Among the step trials, a randomly chosen target step delay (**TSD**), from a total of three (16 ms, 83 ms, 150 ms), separated the two stimuli. The random interlacing of step and no-step trials and the TSDs ensured that the trial type could not be predicted.

### Behavioral evidence of parallel programming of saccades

The actual time available for parallel programming (*parallel processing time* **PPT**; equivalent to delay **D** of Becker and Jürgens, 1979) is the time when both saccade plans are underway, i.e., the time period from the appearance of the second target, to the end of the first saccade. If saccade programming were strictly serial, i.e. the planning of the second saccade starts only after the first is completed, then irrespective of the time available for parallel programming (long or short PPT), the ISI would be invariant (**Fig. 2A**). On the other hand, if the second saccade can be programmed in parallel with the first, then longer the PPT, the greater is the degree of parallel programing possible, resulting in the second plan not having to ‘wait’ for the first to finish, giving rise to shorter ISIs (**Fig. 2B**). Since the ISIs decrease with PPT in the parallel programming condition, the ISI-PPT plot would have a negative slope. The relation between PPT and ISI – specifically the *slope* of the ISI-PPT plot – serves as a metric for the concurrent programming of saccades (Ray et al. 2004).

**Figure 2.**
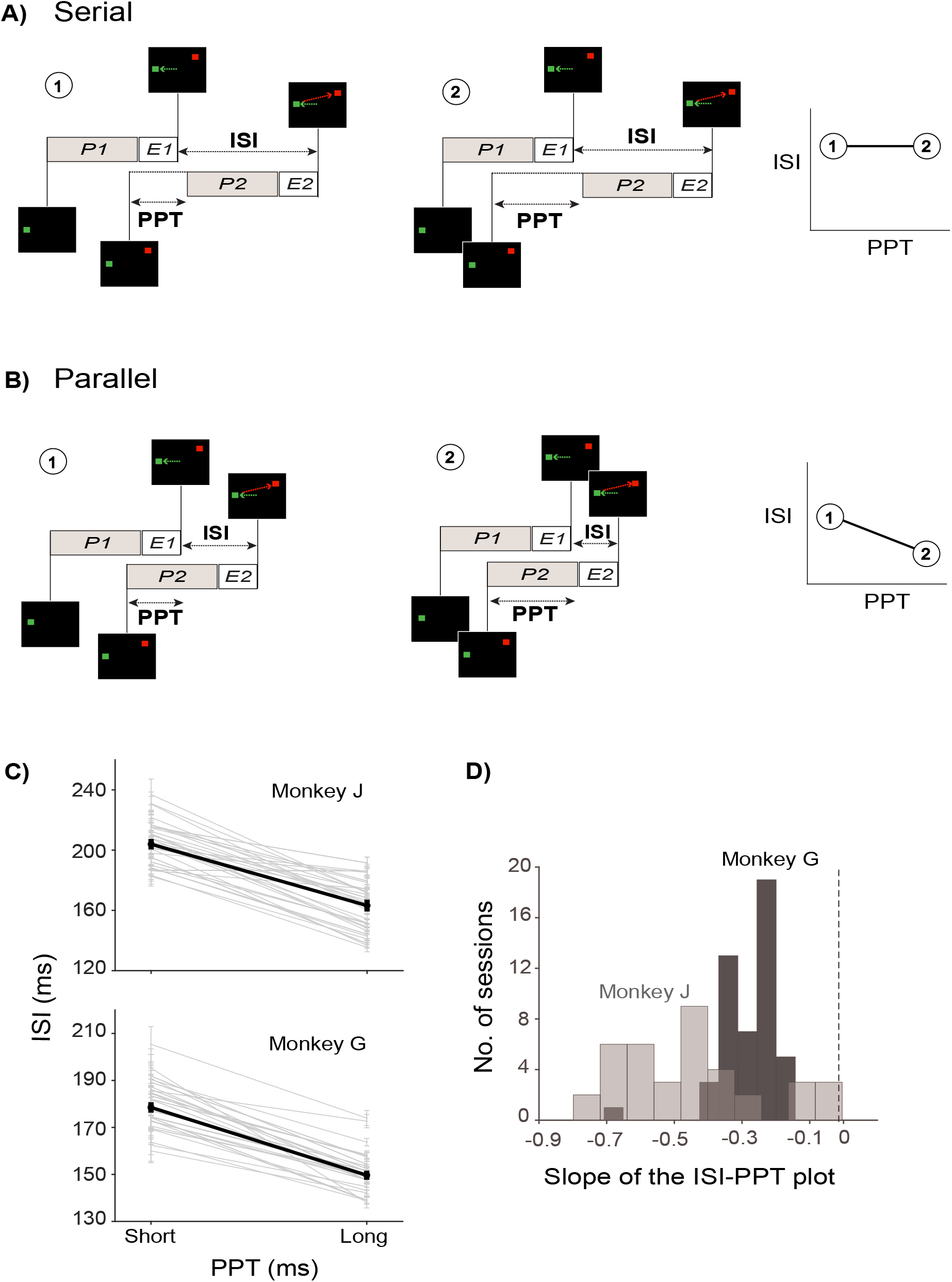
Behavioral evidence of parallel programming. **A**. Each composite saccade program has been divided into a planning stage, and a movement execution stage, marked by ‘*P*’ and ‘*E*’, respectively. (***1***) depicts the condition where the second target comes much later in time after the first, leading to a shorter *time available* for parallel programming (***PPT***). When the second target comes right after the first (***2***), the PPT is longer. In the serial processing condition, the second plan can only start after the first saccade is completed, thus the inter-saccadic interval (***ISI***) remains constant irrespective of the PPT. The slope of the ISI-PPT graph (*right panel*) tends towards 0. **B**. In case of parallel programming of sequential saccade plans, as the PPT increases, the second plan starts earlier in time relative to the first. This results in a decrease of ISI with PPT, and the slope of the ISI-PPT plot becomes negative. **C**. ISI-PPT plots for monkeys J & G across all recording sessions. ISI decreases significantly with PPT. The means and associated standard error bars are shown in bold black lines. **D**. Histograms of slopes of the ISI-PPT plots for individual sessions, for monkeys J (*light fill*) & G (*dark fill*) across all recording sessions.

Step-trials were divided into 2 PPT groups (*short: 0-150 ms and long: 150-300 ms*) and for each session, the mean ISIs for the two PPT groups were calculated. Since ISIs longer than typical saccadic reaction times unambiguously point towards serial processing, trials with inter-saccadic intervals more than the 95^th^ percentile of the latency of no-step saccades (~ 300 ms) were excluded from the analysis. This restricted the trials to those in which parallel processing could have been employed, allowing for better evaluation of whether the mode of saccade sequencing was serial or parallel. At the population level, ISIs for the low and high PPT groups were significantly different for both monkeys (**Fig. 2C**; ANOVA, *F_J_* (1, 74) = 123.75, p = 1.86*10^−17^; *F_G_* (1, 94) = 229.45, p = 5.79*10^−27^). Fig. 2D shows that the slopes of the ISI-PPT plot obtained from individual sessions were significantly negative for both the monkeys (two-sided Wilcoxon signed rank test, *Z_J_* = −5.37, p = 7.71*10^−8^; *Z_G_* = −6.03, p = 1.62*10^−9^), indicating that the ISI decreased with increase in PPT, consistent with the prediction of parallel programming. The negative slopes of the ISI-PPT plot corroborate with previously published behavioral evidence of parallel programming of sequential saccades.

### Classification of FEF neurons

The central goal of this study was to determine if the activity of FEF movement neurons show evidence of concurrent saccade planning. To this end, neural activity was recorded from 163 FEF neurons from two monkeys as they performed the sequential saccade task. Visual (V), visuomovement (VM), and movement (M) neurons were categorized using the memory-guided saccade task according to established criteria (Bruce and Goldberg 1985; Segraves and Goldberg 1987). Neuronal activity was aligned either on target onset (**Fig. 3**, left) or on saccade onset (**Fig. 3**, right) and the activity within the response field (RF; blue traces) was compared to the activity outside the RF (black traces). Following the conventional classification scheme, movement neurons were identified by a presaccadic ramp-up of activity (**Fig. 3A**),visual neurons were identified by a transient burst linked to the stimulus alone (**Fig. 3C**), and visuomovement neurons by the presence of both (**Fig. 3B**). A visuo-motor index (VMI) was calculated for each neuron as VMI = (VA − MA)/(VA + MA) to quantify the relative proportion of visual and movement activity (VA: visual activity, MA: movement activity; see Materials and Methods for details).Thus, positive values indicate the dominance of visually-related activity over movement-related activity and negative values reflect the opposite. The final cohort for subsequent analysis had 84 selected cells: 38 movement neurons (18 from monkey J, 20 from monkey G) whose VMI values ranged from −.97 to −0.32 (mean = −0.7), 34 visuomovement neurons (15 from J, 19 from G) whose VMI values ranged from 0.33 to −0.67 (mean = −0.096), and 12 visual neurons (6 from J, 6 from G) whose VMI values ranged from 0.49 to 0.96 (mean = 0.67).

**Figure 3.**
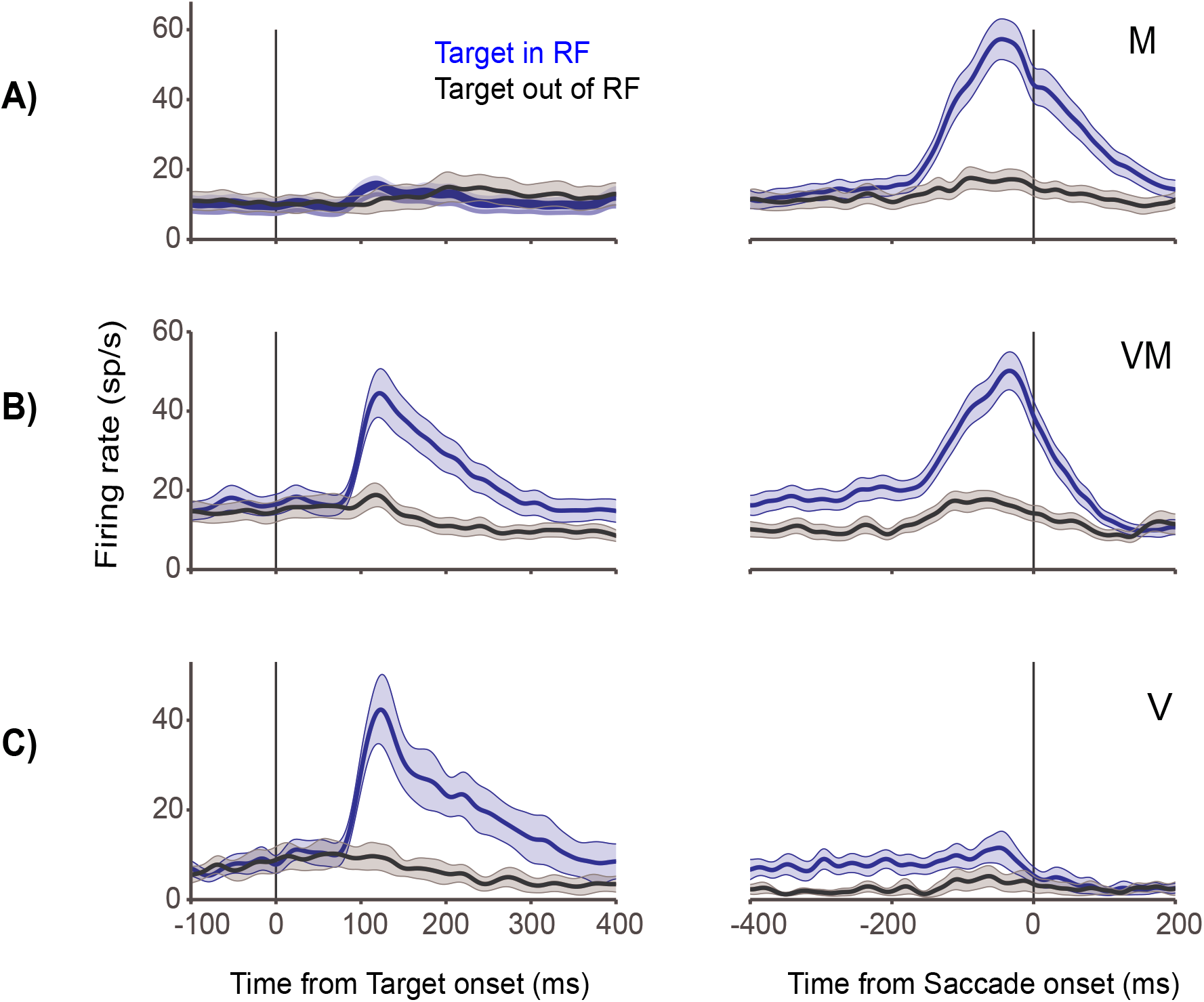
Classification of FEF neurons. Neural activity during the memory-guided saccade task. Traces show the mean firing rate for memory-guided saccades into (*blue*) or out of (*black*) the response field, with activity aligned either target onset (*left*) or saccade onset (*right*). Shaded areas indicate 1 SEM. FEF neurons were classified as Movement (M), Visual (V) or Visuomovement (VM) based on the activity profiles. (**A**), (**B**), (**C**) show average activity profiles for 38 M, 37 VM, and 12 V neurons, respectively.

### Concurrent activity of FEF movement neurons for the second saccade plan

In the context of the FOLLOW task, the critical measure is the onset of activity build-up specific for each saccade plan, i.e. the **neural selection time (NST)**. If sequential saccades are programmed serially, then the NST for the second saccade can be only be after the first saccade has landed and after visual feedback of the new eye-position reaches the frontal areas. The earliest visual feedback delay to FEF is about 50 ms (Schmolesky et al. 1998; Pouget 2005) and therefore, for serially planned saccades, movement neuron activity for the second saccade would only be able to rise 50 ms after the end of the first saccade (**Fig. 4A**). It can be predicted thus, that for saccades planned in parallel, the second saccade NST would occur at least before visual feedback is registered following the first saccade (**Fig. 4A**). A second prediction, based on the behavioral framework put forth in **Fig. 2B**, would be that as the parallel processing time or PPT decreases, the NST would shift later in time (**Fig. 4B**), with respect to the end of the first saccade. Thus, using these two predictions, the NST was used as a distinguishing criterion to compare serial or parallel activation of FEF movement neuron activity.

**Figure 4.**
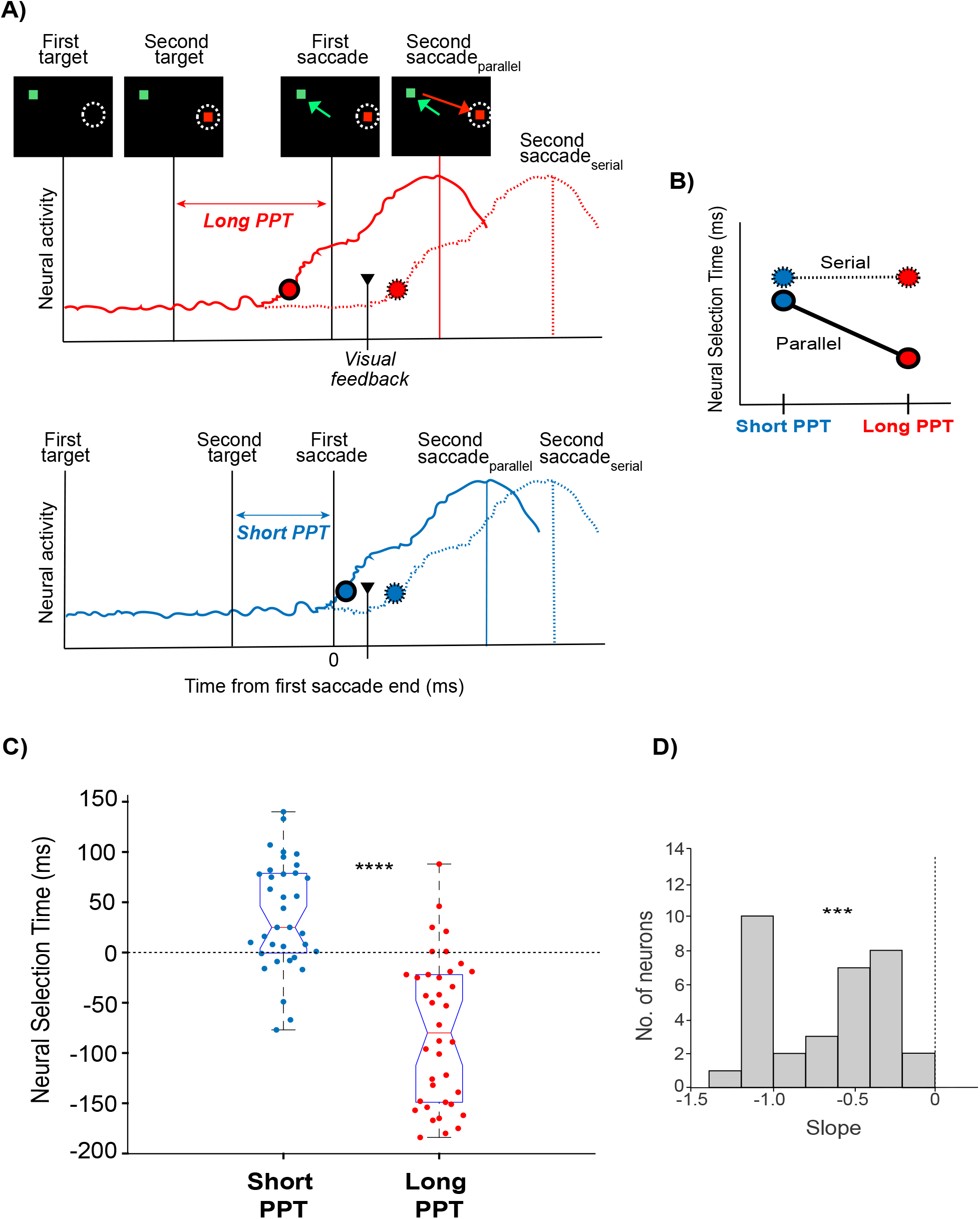
Activity of FEF movement neurons shows evidence of parallel programming. **A**. Schematic showing hypothesized FEF movement neuron activity under short PPT (*blue*) and long PPT (*red*) conditions. The activity is aligned to the end of the first saccade. Task events are shown at the top for *RFin* step trials, where the second saccade was made into response field (*dotted white circle*). Filled red and blue circles mark the neural selection time (NST), or the onset of presaccadic activity specific for the second saccade. In the long PPT condition (*top panel*), activity for serially planned second saccades can only start after visual feedback of the first saccade end reaches the FEF (*red filled circle with dashed border*). If parallel programming is actuated by the FEF movement neurons, then the NST will be much earlier, possibly even before the start of the first saccade (*red filled circle with solid border*). *Bottom panel:* For the short PPT condition, the NST for serially programmed second saccades will be invariant from the long PPT condition, as in both cases, the activity starts much after the first saccade has landed. In the parallel planning condition however, shortening the time available for parallel programming will shift the NST later in time with respect to the first saccade end. **B**. The NSTs derived from the schematic in (**A**) are plotted against the PPT, contrasting the modulation of NSTs with short and long PPT for the serial, and parallel planning conditions. Similar to the ISI-PPT plot in **Fig.2**, the slope of the NST-PPT graph is hypothesized to the negative in the condition where two saccades are programmed in parallel. **C**. Boxplots show neural selection times across the FEF movement neuron population (n = 38). The NSTs showed an inverse relationship with PPT, decreasing significantly from the short to the long PPT group. The horizontal dotted line indicates the first saccade end. In the long PPT group, the NSTs occurred even before the first saccade was initiated. **D**. The slopes of the NST-PPT plots for each movement neuron are shown; the slopes were significantly < 0.

Trials in which the second saccade was made into the RF (*RFin trials*, **Fig. 4A**; see Materials and Methods for details) were selected so that activity specific for the second saccade can be analyzed for short and long PPT groups. The composite neural activity in the *RFin* trials might contain a non-negligible component of *outside-RF* activity from the first saccade. To factor that out, the difference between *RFin* activity and latency-matched no-step activity occurring during saccades made to positions outside the RF was taken. The NST was determined as the time when the difference histogram significantly crossed baseline level, marking the point when activity distinct for the second saccade emerged (see Materials and Methods for details). **Fig. 4C** shows that across the FEF movement neuron population, the NSTs or the onsets of activity related to the second saccade occurred before feedback delays for the short PPT group, and shifted significantly earlier in time for the long PPT group, thus validating the predictions and constituting neural evidence of parallel programming (Kruskal-Wallis, χ^2^ (1, 71) = 34, p = 5.52*10^−09^). The majority (22/38) of the movement neurons selected for the second saccade in the long PPT condition even before the first saccade had started, and 35/38 neurons responded for the second saccade before the first saccade was terminated. While the neural selection time occurred later in the short PPT group, 9/38 neurons still modulated their activity before the first saccade ended. All the neurons showed the inverse NST-PPT relation and exhibited negative slopes (**Fig. 4D**). At the population level, the average slope was significantly < 0 (two-sided Wilcoxon signed-rank test, *Z* = −5.09, p = 3.50*10^−9^). To rule out the possibility of saccadic latency differences to be the sole cause of the observed NST-PPT modulation, a subset of *RFin* trials, where RT was matched across the PPT groups, was taken for every movement neuron. The inverse NST-PPT relation was maintained even with latency-matched trials, with the average slope being significantly less than 0 (two-sided Wilcoxon signed-rank test, *Z* = −2.51, p = 0.0034).

### Parallel programming in FEF visual-salience neurons

The modulation of the onset of activity encoding the second saccade, relative to the time of the first saccade, provides the main neural evidence of parallel programming of saccades in this study. Having delineated and assessed the evidence from the movement neuron population, a similar NST-PPT analysis was done for the visual and visuomovement neuron population. Previous evidence has shown that target selection (undertaken by visual neurons) is distinct and temporally ahead of response preparation (undertaken by movement neurons; Thompson et al. 1996). It logically follows that if evidence of parallel programming can be observed at the level of movement neurons, it should be observable in the activity of the visually-related neurons as well. Consistent with this idea, NSTs for the population of visual and visuomovement neurons shifted earlier in time as the parallel programming time or PPT increased. The effect was significant both at the population level (Kruskal-Wallis, ***V***: χ^2^ (1, 22) = 15.01, p = 0.0001; ***VM***: χ^2^ (1, 68) = 34.06, p = 6.81*10^−12^), and at the individual neuron level, wherein negative NST-PPT slopes were obtained for every neuron (two-sided Wilcoxon signed rank test ***V***: p = 0.002; ***VM***: Z= −5.06, p = 3.9* 10^−5^).

**Fig. 5** shows the cumulative distributions of the neural selection times of FEF movement, visuomovement, and visual neurons. This figure validates the two predictions put forth regarding neural correlates of parallel programming: first, the activity encoding the second saccade emerged before the end of the first saccade for most of the neurons, suggesting that two saccade plans can be co-active. This is observed especially in the long PPT group (Median NST_Visual_ = −150 ms; Median NST_VisMov_ = −102 ms; Median NST_Mov_ = −80 ms), for all the three neuronal populations. Additionally, as the PPT or the time available for parallel programming decreased, the NSTs of all the three types of neurons shifted later in time—this parametric modulation provided further neural evidence of parallel programming. Finally, within each PPT group, the NST showed temporal ordering: Median NST_Visual_ (−15 ms) < Median NST_Vismov_ < (20 ms) Median NST_Mov_ (52 ms) for the short PPT group (long PPT values given above), reflecting the visual-to-motor transformation that occurs as part of a saccade plan.

**Figure 5.**
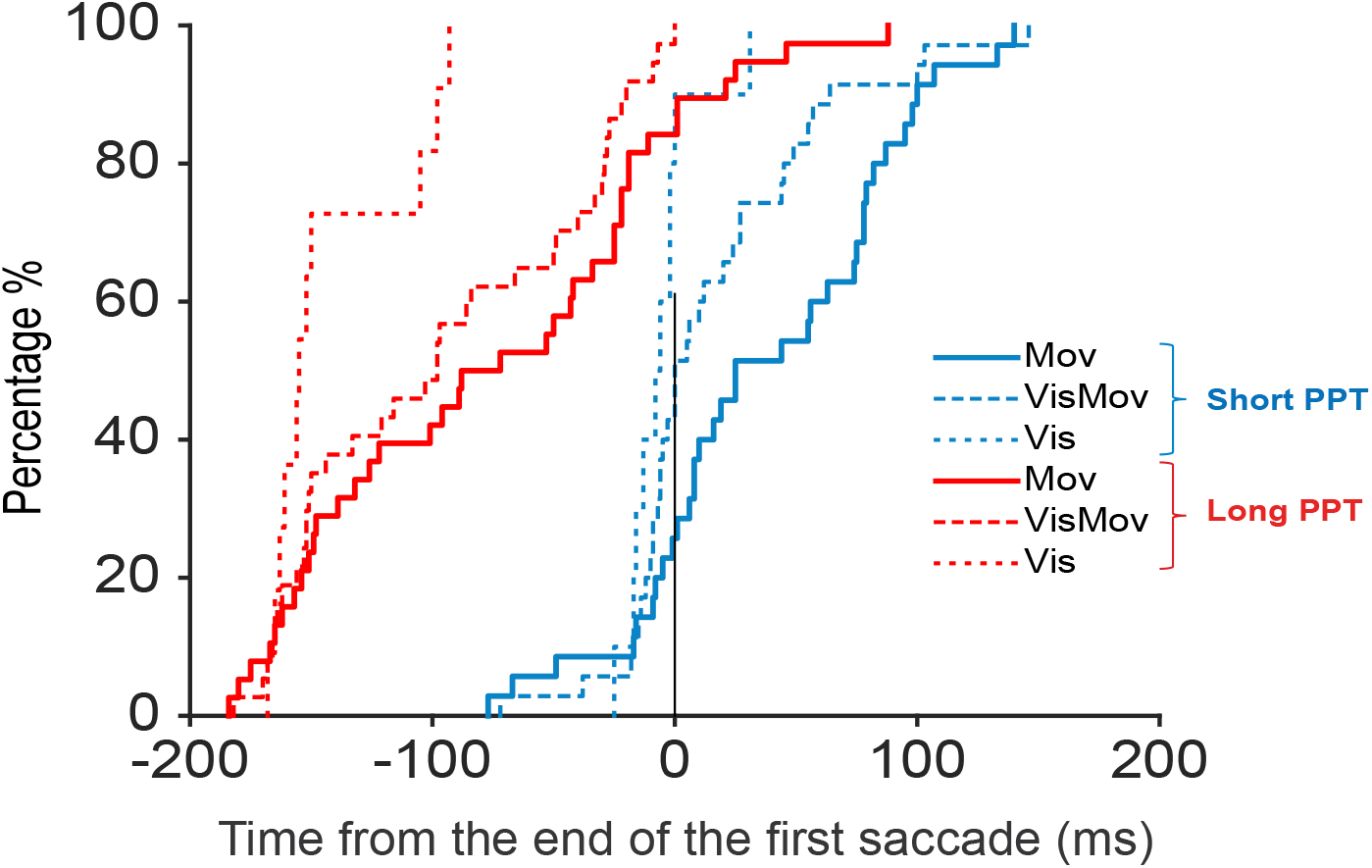
Comparison of neural selection times of visual-salience neurons and movement neurons of the FEF. Cumulative NST distributions for movement (n = 38), visuomovement (n = 34), and visual (n = 12) neurons show that as the PPT is increased from short to long, NSTs across the neuronal population shift earlier, consistent with the predictions of parallel programming. Across all three populations, neural selection times in the long PPT group occurred much before the onset of the first saccade. Within each PPT group, the median NST for visual neurons, preceded that of the visuomovement neurons, which in turn occurred earlier than the median NST for the movement neuron population.

### Vectorial representation of concurrent activity across FEF neurons

The results obtained show that FEF movement neurons could encode future saccade-related activity, even while the eyes were at central fixation and the first saccade plan was underway, in the long PPT trials. During central fixation in the FOLLOW task, the retinotopic vector for the second target is different from the desired saccade vector from the first to the second target. **Fig. 6A** illustrates that for any two targets, accurate generation of the second saccade, the initial *retinal vector 2* must be *remapped* into the final *saccade vector 2*\* (Goldberg and Bruce 1990; Umeno and Goldberg 1997). The question thus arises that whether the early predictive activity, especially in long PPT trials, represents the initial retinal vector of the second saccade goal, or the final remapped second saccade vector. While both cases arise from concurrent programming, the difference lies in the nature of the motor plans encoded. In one case, the first saccade is programmed concurrently with the initial retinal representation of the saccade vector, which indicates that degree of parallel programming is limited, and that much of the processing of the second saccade plan actually occurs after the first saccade is completed. In the second case, all stages of the second saccade plan can proceed in parallel with the first, whether it is reference frame transformation and consequent saccade preparation, without halting at any specific processing step.

**Figure 6.**
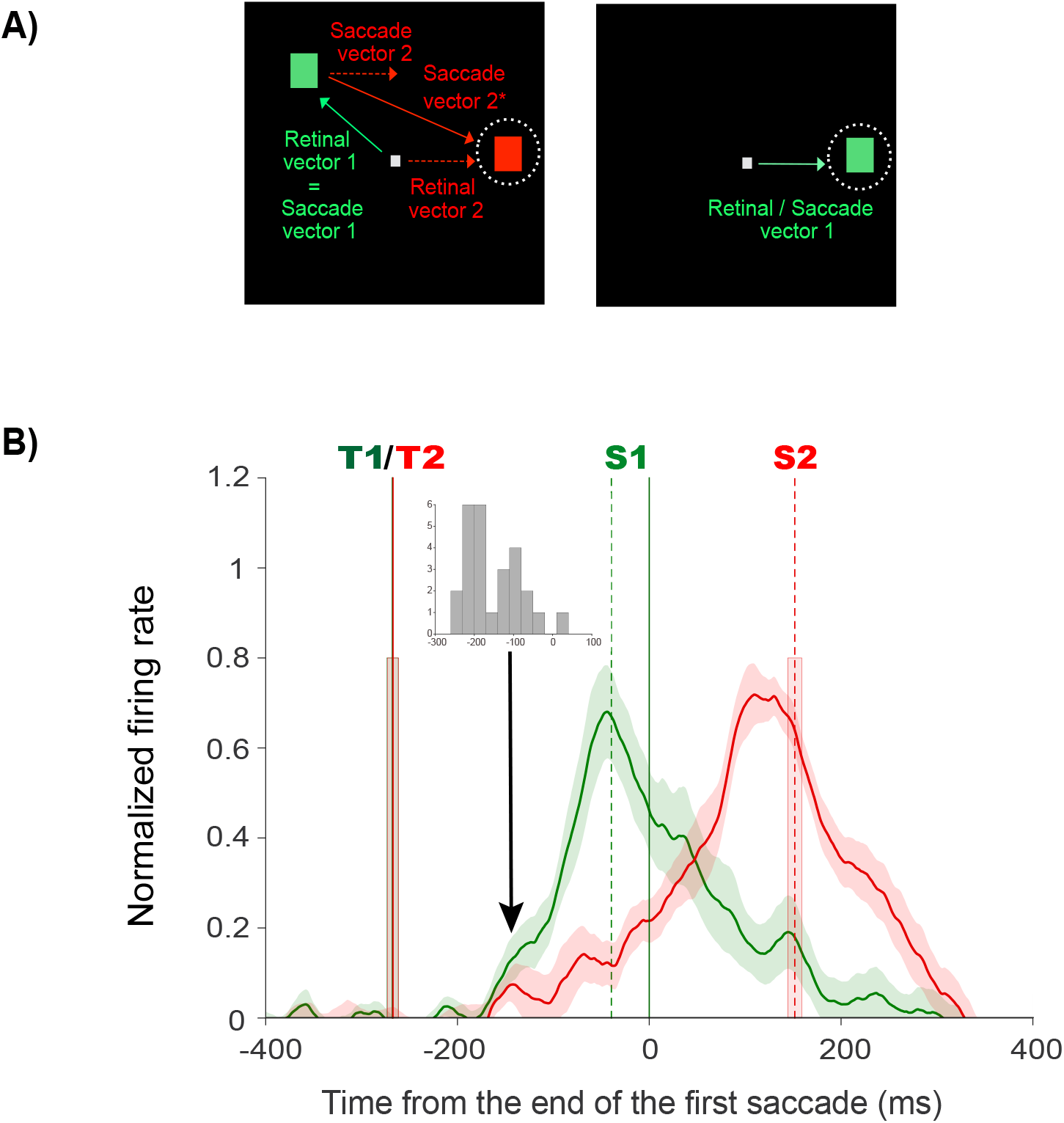
Concurrent movement neuron activity and saccade remapping. **A**. *Left*: In *RFin* trials (second target in response field), at central fixation, the retinal vector of the first target matches with its saccade vector (*green arrow*). However, for the second target, the retinal vector (*red dotted arrow*) has to be remapped to the final saccade vector (Saccade vector 2*; *red solid arrow*) to correctly reach the second target from the first. *Right:* The saccade vector of nostep trials in which the target was in the response field (*white dotted circle*). **B**. Comparison of no-step (*green trace*) and step activity (*red trace*) profiles for the FEF movement neuron population (n = 26). Trials with PPT (S1-T2) in step trials matched to no-step saccade latency (S1-T1) were selected, and neuronal activity was aligned to the first saccade end. The step SDFs diverged from the no-step SDF much before the end of the first saccade, as shown by the *black vertical arrow*. Vertical lines represent mean task markers for no-step (*green*) and step (*red*) trials. The *green dotted vertical line* marks the first saccade onset. *Inset* contains a histogram of the individual divergence points. The downward vertical arrow indicates the mean of the divergence points, with reference to the time on the x-axis.

Under the assumption that the planning of similar vectors would give rise to similar activity patterns, the activity of FEF movement neurons in RFin trials, when the second target was in the RF, was compared to the activity in no-step trials to the same target position (**Fig. 6A**). For each neuron, only one combination of the first and second target positions, with the second target being in the RF, was taken to ensure that the initial and final saccade vectors for the second target remain widely separated. The rationale for this analysis was that since the initial retinotopic vector for the second target was identical in amplitude and direction to the no-step saccade vector, the SDFs for the step and no-step trials would show significant overlap, if the initial predictive activity was coding for the retinal vector of the second target. On the other hand, if the two SDFs diverge out before the termination of the first saccade, it might suggest that the stages further ahead in the second saccade plan, including the generation of the remapped second saccade vector, can be programmed concurrently. The *divergence point* of the spike density functions (SDFs) was calculated in a manner similar to the NST mentioned in the previous section. For accurate comparison of the step and no-step trials, the no-step saccade latency (*S1-T1*) across the population was matched with the PPT (*S1-T2*) of the step trials, as illustrated in the right panel of **Fig. 6A**.

**Fig. 6B** shows the neuronal population data (n = 26), aligned to the end of the first saccade. There is little overlap between the no-step and step SDFs, and the average divergence point (−150.12 ms) occurred much before the first saccade end. The lack of initial overlap suggests that the saccade vector encoding is distinct, and that *RFin* activity in the step-trials represents the remapped *saccade vector 2*\*, and not the *retinal vector 2*. Notably, not only are the SDFs disparate, the onset of movement activity related to the second saccade starts before termination of the first saccade, indicating that *saccade vector 2*\* planning might be able to occur in parallel with the first saccade plan. The inset in **Fig. 6B** shows the histogram of individual neuronal divergence points. Across the movement neurons, the no-step and step-related activity diverged before the first saccade was completed (divergence points < 0; two-sided Wilcoxon signed rank test, *Z* = −4.41, p = 1.05*10^−5^).

Analysis of activity related to retinotopic or remapped vectors suggested that movement neurons can encode the updated second saccade vector in parallel with the first (**Fig. 6B**). A similar analysis, but with increased stringency criteria (see Methods) was done for the visual and visuomovement population. **Fig. 7** shows the neuronal population data for visual (n = 7) and visuomovement neurons (n = 31) aligned to the end of the first saccade. The divergence points for the averaged neuronal activity (V = 9; VM = −92; M = −146) indicates that FEF visual-related cells have a more retinotopic representation of saccade vector, at least in the initial phase, when both saccade plans are underway (i.e. before the end of the first saccade). Movement neurons, on the other hand seem to encode the remapped saccade vector. Thus, activity in the FEF for future saccade has differential vectorial representation based on the neuronal type, and shifts from a retinotopic to remapped encoding from visual to movement neurons. Visually-guided movements necessitate the transformation of visual representations of target location into coordinates appropriate for movement. In case of multiple targets, the movement coordinates have to be updated every time there is an eye movement—parallel programming entails predictive updating of movement consequences to simultaneously plan the future saccade. Given this framework, our results indicate that the visual-to-motor transformation for each saccadic movement is synced with the transformation in vectorial representation from retinotopic-to-remapped coordinates. Our results corroborate with recent evidence showing dynamic sensory-to-motor transitions in the FEF, with movement neurons completing the target-to-gaze coordinate transformation while visual neurons encoded the initial target location (Sajad et al. 2015, 2016).

**Figure 7.**
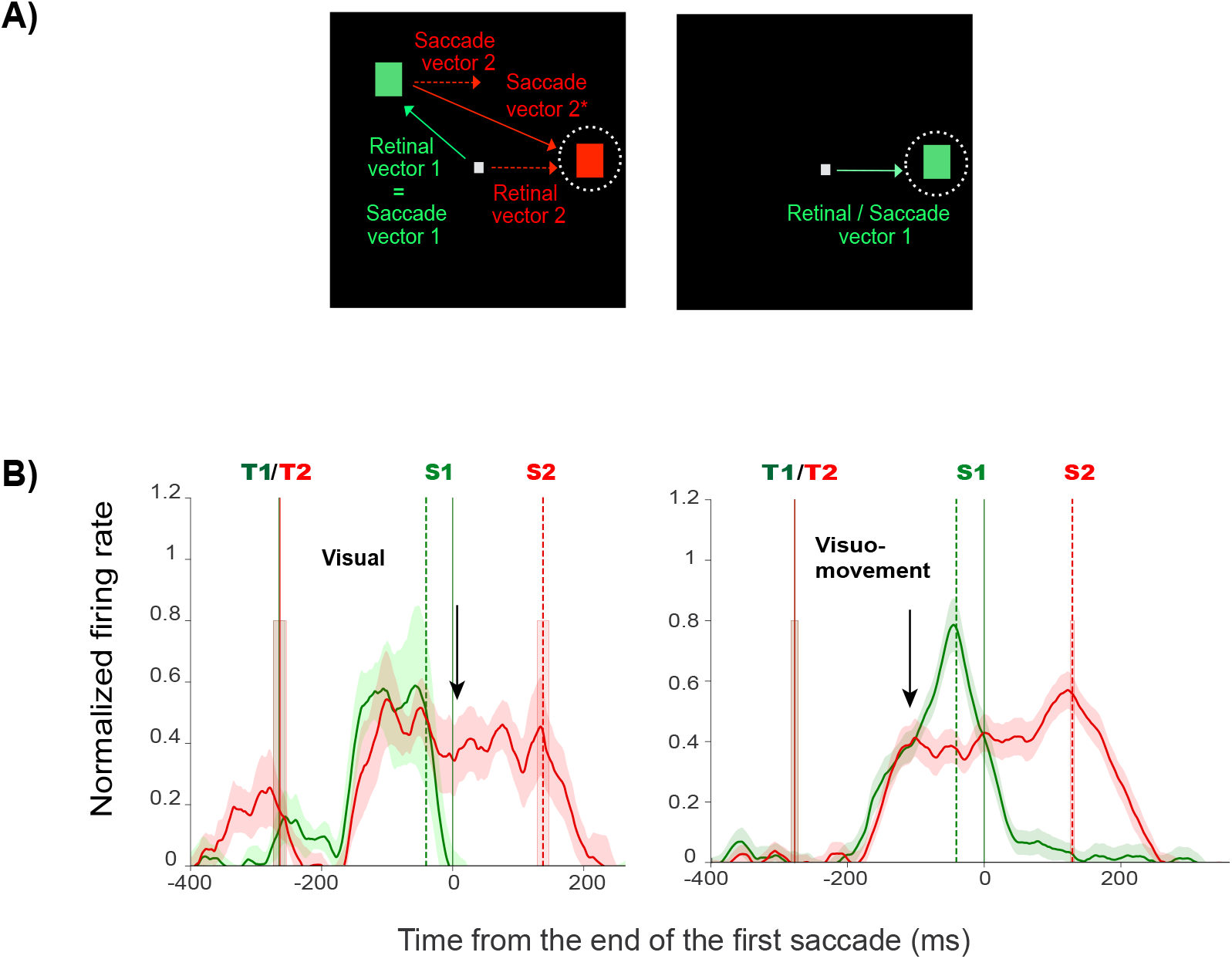
Vector representation in FEF visual-salience neurons. **A**. *Left*: Similar to **Fig. 6A**, showing possible vectorial representations in *RFin* trials (second target in response field) while the eyes were at central fixation. Retinal vector of the first target matches with its saccade vector (*green arrow*). Retinal vector for second target (*red dotted arrow*) has to be remapped to the final saccade vector (Saccade vector 2*; *red solid arrow*) to correctly reach the second target from the first. *Right*: The saccade vector of no-step trials in which the target was in the response field (*white dotted circle*). **B**. Comparison of no-step (*green trace*) and step activity (*red trace*) profiles for the FEF visual (*left*; n = 7) and visuomovement (*right*; n = 31) neuron population. The divergence point for each neuronal type was found from the averaged population activity across neurons and is marked by *black vertical arrows*. Vertical lines represent mean task markers for no-step (*green*) and step (*red*) trials. The *green dotted vertical line* marks the first saccade onset.

## Discussion

This study explored the role of FEF movement neurons in saccade sequencing. The results show that in conjunction with behavioral evidence of parallel programming, neuronal responses related to two sequential saccades can be co-active: movement neurons can become active for a future saccade, while the first one is still being processed or executed. Furthermore, the prospective-saccade related activity started earlier or later in proportion to the actual time available for the oculomotor system to program the two saccades in parallel. The activity of movement neurons map directly onto reaction times, thus our results correspond well with much of the behavioral evidence of concurrent processing of saccades, which comes from temporal indicators. The ability of FEF movement neurons to co-activate multiple saccade plans suggests that predictive activity in the FEF is indexed by the functionally downstream movement neurons (Juan et al. 2004; Purcell et al. 2012), along with visual-salience neurons in the FEF (Phillips and Segraves 2010). In summary, these results constitute the first clear evidence of parallel programming of saccades at the level of FEF movement neurons.

### Concurrent processing of saccade sequences

One assumption in ascertaining the parallel programming of saccades is that each saccade in the sequence is programmed independently, as separate motor plans. However, several behavioral studies on sequence learning have suggested that movement sequences are performed as if they consisted of smaller units of movements or “chunks” (Sternberg 1969; Koch and Hoffmann 2000; Verwey 2001). One prediction of the chunking hypothesis is that the RT of the first movement would be longer for longer sequences (Zingale and Kowler 1987) to account for advance preparation of all the movements in the entire sequence or chunk, grouped together as a single unit. The first saccade RT (mean ± SEM: 264 ± 8.12 ms) in the step-trials of the FOLLOW task used in the current study, was significantly longer (one-way ANOVA, *F* (1, 17) = 9.56, p = 1.25* 10^−3^) than the single saccade no-step RT (mean ± SEM: 221 ± 5.43 ms), indicating that chunking might play a role in pre-planning the saccade sequence. Chunking represents an efficient strategy to weave together stereotyped movements, but it can be argued that chunking represents a form of parallel programming, wherein the entire sequence is being planned at fixation before the first saccade *as one cognitive element*, and does not, per se, change the validity of our results.

Although chunking may provide an explanation for the strong evidence of parallel programming we obtained, the design of the FOLLOW task did not allow for easy, predictable chunking of sequences. First, the two saccade targets in the FOLLOW task always had an angular separation of at least 90° and were present in opposite hemifields, making perceptual chunking difficult. Second, the actual target positions were varied randomly and thus the exact spatial locations of the sequential targets could not be predicted. Third, the temporal delay between the first and the second saccade targets was varied randomly, making temporal predictions about when the second target would come after the first, difficult. Fourth, the sequential-saccade step trials were randomly interspersed with single saccade no-step trials in 30% of the total trials in a session, adding more variability to the task structure. Thus, while the results do not completely rule out chunking as a strategy to stitch together saccade sequences, the task structure accentuates the need to develop two separate saccade plans on every step trial to execute a correct response.

### Parallel programming and rapid error correction

Notably, one other study (Murthy et al. 2007) has looked at FEF movement-related activity in the context of saccade sequences. However, the focus of the study was the role of FEF in rapid error correction; thus, the sequences consisted of an error saccade followed by a corrective saccade. The study found that the movement activity for corrective saccades began before visual feedback of the error was registered. This provided neural evidence of how corrective saccades can be produced faster than reafferent visual processing, suggestive of parallel procesing in error correction. The study used the search-step and the classical double-step tasks, both of which reinforced single saccades, and the two-saccade sequence was an unrewarded consequence of an error response. In contrast, our study reports clear neural evidence of concurrent planning even prior to the execution of the initial saccade, much before registration of visual feedback.

The importance of cognitive context on saccadic planning has been previously established by Ray et al. (2004), who showed that concurrent processing was facilitated in the case of sequences of error and corrective saccades (REDIRECT task), compared to sequences of two correct saccades (FOLLOW task). It has been hypothesized that an error correction system might facilitate the rapid generation of corrective saccades (Schall et al. 2002b, Sharika et al. 2009). However, since the planning of the corrective saccade is only expected to begin once the system has been able to detect the occurrence of the error, or the likelihood of error, (Sharika et al., 2008), it could result in the less robust expression of parallel programming seen in the study by Murthy et al. (2009), as compared to the results observed in this study using the FOLLOW task.

### Predictive remapping of saccade vectors

Computational models of motor control prior to visual feedback, hinge on feedforward control, where the spatial location of the saccade target is compared with the anticipated position of the eyes following the saccade, possibly using an efference copy mechanism in conjunction with an internal forward model (Desmurget and Grafton 2000; Vaziri 2006).Therefore, any movement planning based on the efference copy system can, in principle, start right after the motor command is issued. Thus, parallel programming of saccades presents an interesting problem of encoding the correct vectorial location of the saccade goal. The remapped location for the second target corresponds to the position this target will have on the retina following the first saccade. The presence of correctly executed, rapid saccade sequences with short ISIs from multiple studies, is testimony to the fact that the oculomotor system is able to successfully map the accurate location of the second target, and is not disrupted by the retinotopic shift brought about by the first saccadic movement.

Many neurons in the primate oculomotor system undertake predictive remapping (Goldberg and Bruce 1990; Duhamel et al. 1992; Walker et al. 1995; Sommer and Wurtz 2002), wherein emergence of presaccadic responses selective for the ‘future receptive field’, the position of the RF after the saccade, have been observed. In the FOLLOW task used in this study, the *RFin* trials present an interesting condition: the early concurrent activity occurring during central fixation in the FEF neurons, can represent either the retinotopically mapped activity or the remapped second saccade vector (**Fig. 6A**). Unlike previous studies, the fact that the second target is mapped to the fixation spot by retinotopic coordinates, allows dissociation of vectorial contributions in the neuronal response. Essentially the saccade end-point and neuronal RF remains the same, with different vectorial connections in the planning stage. Our results indicate the possibility of movement-related neurons encoding the accurate, remapped second saccade vector in parallel with the ongoing first saccade plan.

One notable point to consider while interpreting the results is that the analysis depends critically on the assumption that activity in step and no-step trials for planning saccades of similar metrics would be similar. This principle is an oversimplification as the context of a single saccade response and a two-saccade response are widely different, and can influence the saccade-related responses. Even so, given the simplicity and low number of variables of the task, the SDFs of planned and executed responses can be expected to harbor similarities, if coding for similar saccade vectors. Further, our results are consistent with those from a recent behavioral study (Bhutani et al. 2017) which showed using an adaptation task that the remapped second saccade vector, can be planned before the first saccade, and proposed that a representation of the ‘prospective or future motor vector’ is present in the brain, independent of the retinotopic target vectors.

Vector analysis of the activity of FEF visual and visuo-movement neurons indicated that the early visual processing was retinotopic with the remapped vector diverging out around the time of transition from visual to motor activity (in visuomovement neurons). Thus, while rapid encoding of the retinotopic goal vector is facilitated by visually-responsive neurons of the FEF, clear evidence of remapping of the motor vector is observed only at the level of movement-related activity. Presaccadic attentional shifts to the remapped location has been shown to decrease saccadic latencies to subsequent targets, providing a tangible functional benefit of predictive remapping (Rolfs et al. 2011). Such functional remapping may be brought about by the FEF movement neurons, aiding concurrent and accurate spatial representation of multiple saccade targets, ultimately leading to a rapid executed sequence of saccades. An analysis of the microcircuitry of remapping in the FEF (Shin and Sommer 2012) yielded that neurons in layer V displayed full remapping properties, as opposed to partial remapping shown by partial remapping shown by layer IV neurons. While the cohort analyzed consisted mainly of visuo-motor neurons for layer V, it is well-established that neurons in the FEF that generate movement-related or fixation-related activity are located in layer V and innervate the superior colliculus and parts of the brain stem saccade generator circuitry (Segraves et al. 1987, 1992; Sommer and Wurtz, 2000; Pouget et al 2009). Thus, we propose based on our results that FEF movement neurons are actively involved in the mechanism of saccade vector remapping.

An alternative to the remapping process is the direct exocentric or allocentric representation of saccade goals with respect to visual landmarks (Sharika et al. 2014). The exocentric system uses an object-to-object representational system, where information about the location of one object is encoded using cues from another object in its proximal external space. We believe that such an exocentric representation is unlikely to mediate the observed parallel programming for two reasons. First, exocentric or allocentric representations have been shown to develop slowly across several hundreds of milliseconds, (Hu and Goodale 2000; Westwood et al. 2000; Zimmermann et al. 2013) that typically also involve having stable landmarks. Such stimulus conditions did not exist in our task--which involved the production of rapid visually-guided saccades in random trials and to random locations. Second, exocentric coding of the second target location would predict that the visual representations to also show evidence of remapping, which we did not observe, as we did for movement related activity.

In conclusion, we assessed the role of FEF movement neurons in saccade sequencing. The results show that FEF movement neurons bring about parallel planning at the neuronal level by driving co-activation of presaccadic activity related to multiple consecutive saccades. The extent of co-activation is also modulated by temporal measures of concurrent programming, suggesting that FEF movement neurons are critically involved in the generation sequential saccade behavior.

## Acknowledgements

We thank S. Sengupta for helping with behavioral training and Dr. A. Gopal P.A. for helping with data collection.

## Author Contributions

A.M. conceived the study, and D.B. collected and analyzed the data, and wrote the manuscript draft. A.M. and D.B. both contributed to the experiment design, surgical procedures, interpretation of data, and editing of the manuscript.

## Glossary

ANOVA: : analysis of variance
FEF: : frontal eye field
IM: : intra-muscular
ISI: : inter-saccadic interval
M: : movement
MA: : movement activity
MG: : memory-guided
NST: : neural selection time
PPT: : parallel processing time
RT: : reaction time
TSD: : target step delay
RF: : response field
RFin: : inside response field
RFout: : outside response field
SD: : standard deviation
SDF: : spike density function
SEM: : standard error or mean
V: : visual
VA: : visual activity
VM: : visuomovement
VMI: : visuo-motor index

